# Audio-tactile and peripersonal space processing around the trunk in human parietal and temporal cortex: an intracranial EEG study

**DOI:** 10.1101/249078

**Authors:** Fosco Bernasconi, Jean-Paul Noel, Hyeong Dong Park, Nathan Faivre, Margitta Seeck, Laurent Spinelli, Karl Schaller, Olaf Blanke, Andrea Serino

## Abstract

Interactions with the environment happen by the medium of the body within one’s peripersonal space (PPS) - the space surrounding the body. Studies in monkey and humans have highlighted a multisensory distributed cortical network representing the PPS. However, electrophysiological evidence for a multisensory encoding of PPS in humans is lacking. Here, we recorded for the first time intracranial electroencephalography (iEEG) in humans while administering tactile stimulation (T) on the trunk, approaching auditory stimuli (A), and the combination of the two (AT). To map PPS, in AT trials, tactile stimulation was delivered when the sound was far, at an intermediate location, or close to the body. We first identified electrodes showing AT multisensory integration (i.e., AT vs. A+T): 19% of the recording electrodes. Among those electrodes, we identified those showing a PPS effect (30% of the AT electrodes), i.e., a modulation of the evoked response to AT stimulation as a function of the distance between the sound and body. For most sites, AT multisensory integration and PPS effects had similar spatiotemporal characteristics, with an early response (~50ms) in the insular cortex, and later responses (~200ms) in pre‐ and post-central gyri. Superior temporal cortex showed a different response pattern with AT multisensory integration at ~100ms without PPS effect. These results, representing the first iEEG delineation of PPS processing in humans, show that PPS processing happens at neural sites where also multisensory integration occurs and at similar time periods, suggesting that PPS representation (around the trunk) is based on a spatial modulation of multisensory integration.

## Introduction

The space immediately adjacent to and surrounding the body - defined as peripersonal space (PPS) (di Pellegrino et al., 1997; Rizzolatti et al., 1981, 1997) - is particularly relevant for behavior, as it is where physical interactions with the environment occur (Graziano and Cooke, 2006; Làdavas and Serino, 2008). The ecological significance of the PPS is evidenced in that the primate brain has developed a fronto-parietal network encoding preferentially multisensory stimuli occurring near to (as opposed to far from) the body. That is, neurons located in monkey posterior parietal cortex (i.e., intra-parietal sulcus (IPS) (Duhamel et al., 1997, 1998)), area 7b (Leinonen and Nyman, 1979a, 1979b), (Fogassi et al., 1996; Graziano et al., 1997), and ventral premotor cortex (vPM; (Fogassi et al., 1996; Graziano et al., 1997) have been reported to respond to both tactile stimuli on body parts, as well as to visual (Schlack et al., 2005) or auditory (Graziano et al., 1999; Schlack et al., 2005) stimuli occurring near the same body part that is touched.

A homologous PPS neural network is postulated to exist in humans, supported by numerous psychophysical (Salomon et al., 2017; Spence et al., 2004) and neuropsychological (Farnè and Làdavas, 2000; Maravita and Iriki, 2004) studies demonstrating enhanced processing of tactile stimulation when a task-irrelevant visual or auditory object is present near vs. far from the body. These studies rely on the congruent presentation of multisensory stimuli in the environment (Serino et al., 2015) and (Canzoneri et al., 2012) are body part centered (hand: (Canzoneri et al., 2012); face: (Teneggi et al., 2013); Trunk: (Galli et al., 2015; Noel et al., 2015, 2015)). The existence of a homologous PPS neural network in humans is further supported by fMRI studies, which have demonstrated a close association between the areas encoding for PPS in non-human primates and humans (e.g., Bremmer et al., 2001; Brozzoli et al., 2011, 2012; Ferri et al., 2016; Gentile et al., 2011; Grivaz et al., 2017; Makin et al., 2009). In addition to the above mentioned PPS areas described in monkeys, human fMRI has equally revealed primary somatosensory cortex (S1), parietal operculum (e.g., Tyll et al., 2013), insula (e.g., Schaefer et al., 2012), cingulate cortex (e.g., Holt et al., 2014) and the lateral occipital cortex (Gentile et al., 2013) as brain regions encoding PPS (for a review see Grivaz et al., 2017).

The characterization of the areas encoding PPS in humans, however, has quasi-exclusively mapped the peri-hand representation (Brozzoli et al., 2011, 2012; Gentile et al., 2011; Makin et al., 2007), with only a few studies investigating the peri-face space (Bremmer et al., 2001; Holt et al., 2014; Sereno and Huang, 2006 for exceptions) and even fewer on the peri-trunk space (see Huang et al., 2012 for an exception). Moreover, while the encoding of PPS is largely taken to be subsumed by multisensory networks, most of the evidence on PPS-related neural response is based on the finding that PPS neurons or regions respond both to tactile and visual (or auditory) stimulation. Yet, only one single electrophysiological study (Avillac et al., 2007) has demonstrated clear multisensory integration (i.e., multisensory supra‐ or sub-additivity, see below) (see Gentile et al., 2011 for fMRI evidence). Finally, evidence on the PPS system in humans mainly comes from fMRI studies, which are limited by the fact that the BOLD response is not only an indirect measure of neural activity but also one that lacks the temporal resolution needed to characterize the dynamics of neural computations leading to the encoding of PPS. Thus, the existing literature has left several open questions such as whether visuo-tactile or audio-tactile PPS processing is truly multisensory and whether multisensory integration and PPS processing occur at similar time periods. Answering these questions would provide insight on whether the spatial modulation of multisensory processing, characterizing PPS representation, occurs in parallel with multisensory integration or follows it in a hierarchical way.

Here, we address the issues raised above, by recording intracranial electrical brain activity in humans, via surgically implanted electrodes in six patients with pharmacoresistant epilepsy. By combining high temporal and spatial resolution, intracranial recordings overcome some of the limitations of the techniques used in previous PPS experiments. Patients received tactile stimuli on the trunk while a task-irrelevant auditory stimulus approached the body. Because of the novelty of the study (and therefore limited evidence to generate hypothesis-driven analysis), and to avoid biases induced by prior assumptions, we used a data-driven methodology. To test multisensory PPS processing, we adopted a 2-step analysis approach, in which we first identified electrodes demonstrating veritable multisensory integration – defined as showing non-linear sensory summation of response to multisensory stimuli (i.e., A+T vs. AT; Stein and Stanford, 2008; Stein et al., 1993) – and then, within the resulting set of multisensory sensors, we search for electrodes showing a neural response that is modulated by the distance between the location of tactile and auditory stimulation (see Quinn et al., 2014 for a similar analytic approach in the visuo-tactile domain). By comparing the sites and the timing of multisensory integration and PPS processing, we investigated whether multisensory brain areas also encode for PPS.

## Methods

### Participants

Intracranial EEG data (i.e., local field potentials; LFP) were recorded from 6 epileptic patients (3 females, 2 left-handed, mean age: 33±4.8 (mean ± sem), see supplementary Table S1 for age, gender, handedness, and epilepsy focus of each patient) who were either implanted stereotactically with depth electrodes and/or grid electrodes were placed on the cortical surface (P-1, P2 and P-5) for clinical purposes (i.e., pre-surgical evaluation in pharmacoresistant epilepsy, see Table S2 for details). Written informed consent was obtained from all patients to take part in the procedures, which were approved by the local ethics committee.

### Material and apparatus

Tactile and auditory stimuli were administered during the task (see Procedures below). Tactile stimulations were applied to the patient’s chest, on the upper part of the sternum, by activation of a vibro-tactile motor (Precision MicroDrives, shaftless vibration motors, model 312-101, 3 V, 60 mA, 9000 rpm, 150 Hz, 5g, 113 mm2 surface area, maximal rotation speed reached in 50ms). Tactile stimulation lasted 100ms and was controlled via a purpose-made microcontroller (Arduino™, http://arduino.cc, refresh rate 10 kHz) and driven by in-house experimental software (ExpyVR, http://lnco.epfl.ch/expyvr, direct serial port communication with microcontroller).

The auditory stimulus consisted of a white noise sound, which was approaching from the front, and centered on the patient’s body, presented via insert earphones (model ER-4P; Etymotic Research). To give the impression that the sound was approaching from the front, sounds were pre-recorded from two arrays of 8 speakers (2m length in total) and head model binaural microphones (Omni Binaural Microphone, http://3diosound.com, see Serino et al., 2015 for detail regarding the external auditory setup).

### Procedures

During the experiment, the patient was comfortably lying in bed, with the upper part of the body reclined forming approximately a 135° angle with the rest of their body. The patient was asked to keep their eyes closed for the duration of the experiment, and they were equally instructed to be attentive to the approaching sound and tactile vibrations. No overt task was requested from the patients.

The experiment consisted in three different types of trials: i) Auditory trials (unisensory audio; A), which consisted of an approaching sound, with a maximal simulated distance from the body of 2m (and lasted a total of 3 seconds; speed: 0.66 m/s), ii) vibro-tactile trials (unisensory tactile; T), which consisted of three successive stimulations, administrated 500ms, 1500ms and 2500m after the onset of the trial, and iii) audio-tactile trials (multisensory; AT), in which the tactile stimulation were administrated 500ms (Far distance, equivalent to 1.7m from the body), 1500ms (Middle distance, equivalent to 1m), and 2500m (Close distance, equivalent to 0.3m) after the initiation of the trial and auditory stimulus onset. To prevent anticipation effects on the vibro-tactile stimulation, a jitter of 0-200ms (step of 50ms) was used for each delay of stimulation. This small temporal jitter allowed us to induce some variability in the timing of tactile stimulation, while not altering the spatial position of the sound when tactile stimulation was administered. A total of 85 trials for each condition were presented, in a randomized manner. The inter-trial interval was shuffled randomly between 1.4s, 1.7s, or 1.9s. In total, the experiment lasted approximately 20 minutes.

### Electrode implantation, intracranial EEG recordings, and pre-processing

In total, 500 electrodes (depth & grid) were implanted in 6 patients, covering diverse cortical and subcortical areas including the post‐ and pre-central gyrus, insula, temporal and parietal operculum, amygdala, hippocampus, frontal and temporal cortex (see Figure 1 for the location of all recording sites). All implantation sites were determined purely based on clinical requirements. Three different types of electrodes were used for the recording: standard electrodes (contact size: 2.4mm, inter-electrode spacing: 10mm) ‘short spacing’ electrodes (contact size: 1.32mm, inter-electrode spacing: 2.2mm), and ‘micro’ electrodes (contact size: 1.6mm, inter-electrode spacing: 5.0mm).

**Figure 1.**
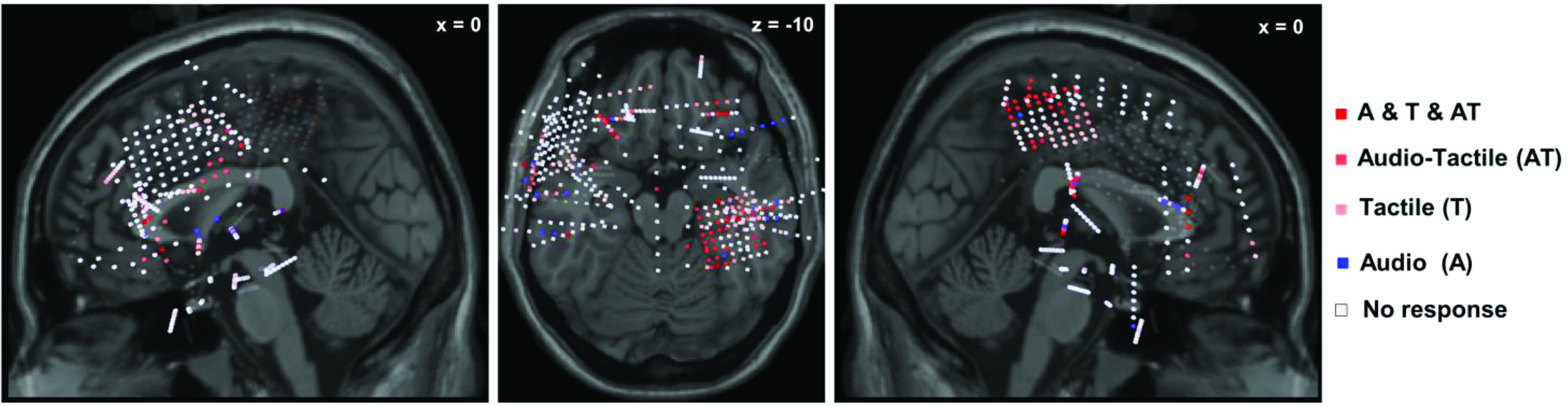
Locations of all recording sites in 3D MNI space. MNI coordinates of electrodes from all 6 patients (500 electrodes in total) plotted on the Colin27 MRI template (sagittal and axial planes). Note that locations are in 3D MNI space, and not located on the surface of MRI slice shown (thus, recording sites behind the depicted MRI slice are marked with faded color). In white, the implanted electrodes not showing a response (vs. baseline, cluster-corrected) to stimuli, in blue, the electrodes showing a response to audio stimuli only, in light red, the electrodes showing a response to tactile stimuli only, in red, the electrodes showing a response to audio-tactile stimuli, and in dark red the combination of the three.

For each patient, intracranial EEG signals were simultaneously recorded across all sites (Micromed System PLUS, Micromed, Mogliano Veneto, Italy) with a sampling rate of 2048 Hz, and an online high-pass filtered at 0.02 Hz. The external reference electrode was located at position Cz (i.e., vertex). Continuous intracranial EEG data were down-sampled to 512 Hz for analyses. Signals were filtered with a band-pass filter between 1Hz and 40Hz. Initial peri-stimulus EEG epochs were generated (800ms pre-trial onset – auditory stimulus in the case of A and AT trials - to 3000ms post-trial onset), and each epoch was centered to zero. Data were further re-epoched to 100ms pre-stimulus onset to 300ms post-stimulus onset. Baseline correction on the 100 ms pre-stimulus onset was applied, only on the electrodes that were identified as responsive vs. baseline (see Statistical analysis below for details).

In each patient, electrodes and trials showing excessive noise (i.e., > 6 interquartile range) were excluded, and thus 480 clean electrodes out of 500 implanted electrodes were used for further analysis. On stripes and depth electrodes, bipolar signals were computed by subtracting intracranial EEG signals from two adjacent electrodes (e.g., A1 – A2, A2 – A3…) from within each electrode shaft, to eliminate the influence of the common external reference and remote sources (Lachaux et al., 2003). In the case of grid electrodes, as bipolar referencing is not suitable (Lachaux et al., 2012), we computed the average of the grid as a reference (i.e., local reference). After preprocessing the number of trials for the tactile conditions was 80.3 ± 1.2 (mean ± sem), 79.7 ± 1.7 (mean ± sem) for the auditory conditions, and 77.5 ± 1.5 (mean ± sem) for the audio-tactile condition. The number of trials retained per condition was not significantly different (F(2,15) = 0.97; p = 0.39).

To compute the Montreal Neurological Institute (MNI) coordinates for each electrode, a post-implant computed tomography (CT) image was co-registered to the normalized preoperative magnetic resonance imaging (MRI) using Cartool Software (Brunet et al., 2011). The midpoint between two depth electrodes was considered as the location of the corresponding bipolar derivation, and for the grid electrodes, the exact position was used. Then, locations of the electrodes were visualized on the Colin27 MRI brain template using the BrainStorm toolbox (Tadel et al., 2011). The anatomical description was assessed using Talairach coordinates (http://talairach.org/; Lancaster et al., 1997, 2000), with a 1mm cube around the coordinates defined above.

### Statistical analysis

According to previous literature in non-human primates as well as humans, PPS is defined as a multisensory effect that is space dependent – (see introduction). Therefore, to identify PPS electrodes, we used a two-step statistical approach. First, we first identified electrodes responding to multisensory stimuli (vs. baseline), and among those electrodes, we investigated which responded in a manner suggesting multisensory integration (see below). Second, among the electrodes showing a multisensory integration effect, we characterized those that had a PPS effect – a multisensory response that is dependent on the proximity of exteroceptive signals (e.g., auditory information) to the body (a similar approach has been previously used in iEEG studies, e.g., (Quinn et al., 2014).

Statistical significance within each electrode was assessed through (temporal) cluster-based permutation statistics (Maris and Oostenveld, 2007) as implemented in the Fieldtrip toolbox (Oostenveld et al., 2011). The advantage of this test is that differences between conditions can be identified without prior assumptions about the temporal distribution of effects. Therefore, being a data-driven approach. The cluster-level statistic was calculated as the maximum sum (maxsum) of the t-values within the cluster. Statistical significance at the cluster level was determined by computing a Monte Carlo estimate of the permutation distribution of cluster statistics, using 5000 resampling of the original data, yielding a distribution of cluster-level statistics under the null hypothesis that any differences between conditions are due to chance. Within a single electrode, a cluster was taken to be significant if it fell outside the 95% confidence interval of the permutation distribution for that electrode. The determination of significant temporal clusters was performed independently for each electrode. This method controlled for false alarms within an electrode across time points.

### Active (unisensory and multisensory) electrodes

To evaluate and select active electrodes for latter between-conditions testing, we applied the cluster-based, nonparametric statistical procedure (see above for details). Electrodes demonstrating a significant response (post-stimulus period 0-300ms) relative to a baseline (-100ms to 0ms) during the post-stimuli onset to A, T and/or AT trials were considered as active electrodes (no baseline correction was applied for this analysis).

### Audio-tactile multisensory integration and PPS electrodes

Among the active AT electrodes, we first selected those showing a response revealing significant multisensory integration (i.e., demonstrating either a supra or subadditivity effect., A+T vs. AT), and then among the electrodes evidencing multisensory integration we investigate which had a “PPS effect” – i.e., a non-linear modulation of tactile response depending on the distance of the sounds from the body. To identify both the “multisensory” and “PPS” electrodes a modified version of the cluster-based, nonparametric statistical procedure outlined by Maris and Oostenveld (2007) was applied. To assess statistically a multisensory integration effect, we applied a cluster-based permutation statistic individually to each electrode (which showed to be active in comparison to baseline), with the contrast AT vs. A+T. To investigate the PPS effect, we first computed the difference AT-A (this approach was chosen to assess whether the PPS response pattern was different from any possible habituation/anticipation effect we may observe in the T condition), and then to assess a statistically PPS effect we applied a cluster-based permutation statistic individually to each electrode (which showed multisensory integration), with the contrast Far vs. Middle vs. Close (one-way ANOVA). Further, to investigate if any anticipation/habituation effect had occurred and could account for the PPS effect, we computed a similar analysis as for the “PPS effect” on the condition in which only the tactile stimulus was presented. Electrodes demonstrating both a multisensory integration and PPS effect but no (or at least with different response pattern) tactile habituation effect can arguably be safely considered electrodes evidencing a multisensory effect that is space-dependent, i.e., putatively recording activities from “PPS brain areas.”

## Results

### Active unisensory and multisensory electrodes

We first investigated which electrodes showed a significant response vs. baseline period, across the 480 implanted electrodes (Figure 1, 480 out of 500 electrodes were included after pre-processing), from all 6 patients. Our data show that 104 electrodes (~22% out of 480) measured a response to the T stimulus, 78 electrodes (16% out of 480) measured a response to the A stimulus, and 104 (~22% out of 480) showed to be responsive to AT stimulation (see Figure 1).

### Location of unisensory and multisensory electrodes

Among the responsive electrode, we assessed the distribution of unisensory (T and A) and multisensory (AT) responses. The electrodes responding to the T stimulus were predominantly located in the postcentral gyrus (PCG), but also in the insular cortex (IC) and inferior parietal gyrus (IFG). The electrodes responding to the A stimulus were predominantly located in the superior and inferior temporal gyrus (STG and ITG). The electrodes responding to the AT stimuli were in the PCG, the precentral gyrus (PreCG), the mid-temporal gyrus and STG, IFG, IC and parahippocampal gyrus (PHG) (Figure 1). Next, to give an overview of the distribution of the electrode location (and quantify the proportion of electrodes), we grouped them into larger brain regions. Our results show that in the frontal areas 22% of the electrodes show a response to A, 48% to T and 31% to AT; in the temporo-insular areas 23% responded to A, 46% to AT and 31% to T; in the parietal areas 5% responded to A, 26% to T, and 52% to AT (for details of the electrodes distribution in the brain see Figure 1, and Figure S1).

### Location of the electrodes showing audio-tactile multisensory integration

Among the responsive AT multisensory electrodes (104), 20 electrodes (~19% of active electrodes) showed significant multisensory integration (i.e., AT vs. A+T). This multisensory integration occurred principally within the PCG (7 electrodes, 35% of the AT multisensory electrodes) and the STG (3 electrodes, 15% of the AT multisensory electrodes). It was also found within the PHG (3 electrodes, 15% of the AT multisensory electrodes), within the Pre-CG (1 electrode, 5% of the AT electrodes), IFG (1 electrode, 5% of the AT electrodes), and the IC (1 electrodes, 5% of the AT electrodes). 4 electrodes were situated in the white matter (Figure 2 and see Table 1 for more details).

**Figure 2.**
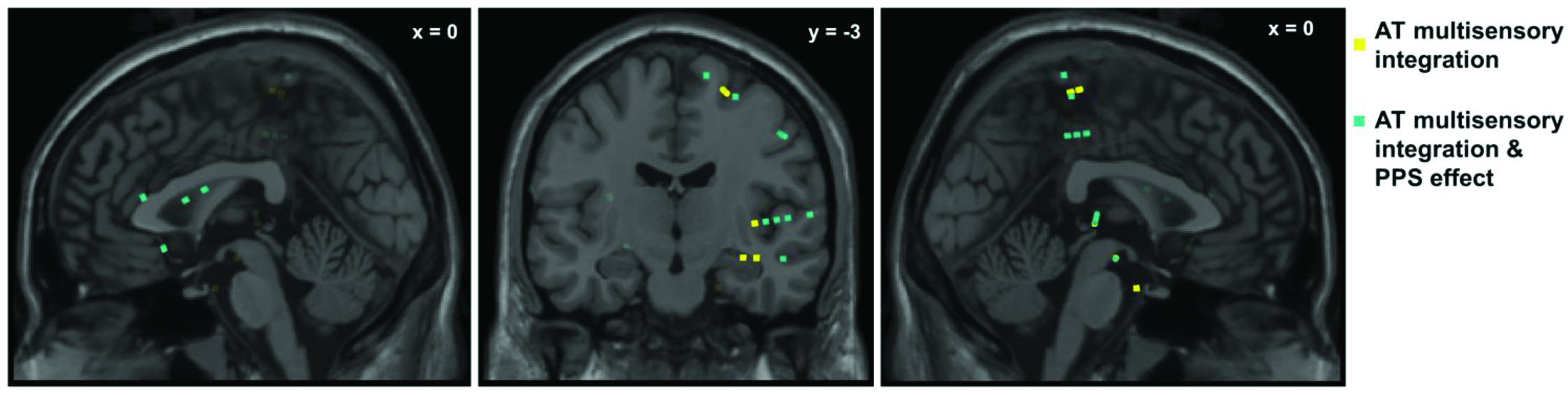
Locations of electrodes showing an AT multisensory integration and peripersonal space (PPS) effect, in 3D MNI space. MNI coordinates of electrodes from all 6 patients, electrodes showing specifically a significant multisensory integration profile are highlighted in yellow (20 electrodes, see Table 1 for the position), electrodes showing both an AT multisensory integration and PPS effect are highlighted in green (6 electrodes, see Table 2 for electrodes positions). Note that locations are in 3D MNI space, and not located on the surface of the MRI slice (thus, recording sites behind the depicted MRI slice are marked with faded color).

**Table 1.**
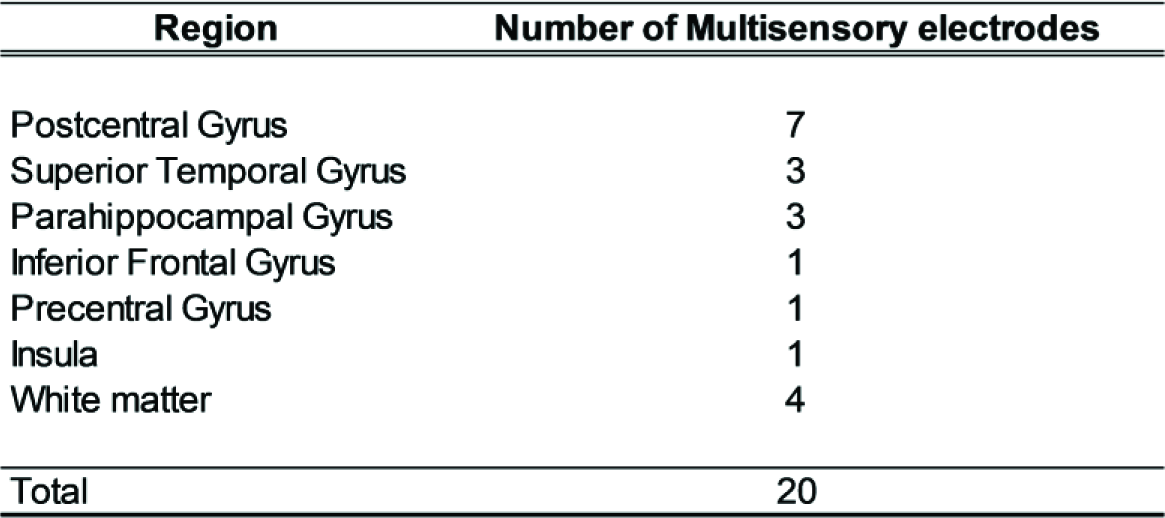
Number of significant electrodes showing AT multisensory integration, sorted by brain regions (p-val < 0.05, cluster-corrected).

| Region | Number of Multisensory electrodes |
| --- | --- |
| Postcentral Gyrus | 7 |
| Superior Temporal Gyrus | 3 |
| Parahippocampal Gyrus | 3 |
| Inferior Frontal Gyrus | 1 |
| Precentral Gyrus | 1 |
| Insula | 1 |
| White matter | 4 |
| Total | 20 |

### Timing of audio-tactile multisensory integration

On average, across the 20 electrodes showing AT multisensory integration, the effect occurred from 151ms ± 18ms (mean ± sem) to 244ms ± 15ms (mean ± sem) post-stimulus onset (i.e., tactile). Within the IC the effect occurred from 63ms to 296ms post-stimulus onset, and a supra-additive non-linear (AT > A+T) neural response was observed. Within the STG the response occurred from 100ms ± 40ms (mean ± sem) to 210ms ± 57ms (mean ± sem), and supra-additive non-linear (AT > A+T; neural response interactions between multisensory and the sum of the constituent unisensory stimuli) was observed. Within the PHG the effect occurred from 112ms± 39ms (mean ± sem) to 239ms ± 25ms (mean ± sem) and a supra-additive non-linear (AT > A+T) neural response was observed between the multisensory response and the sum of the constituent unisensory stimuli. Within the pre‐ and post-central gyri and IFG the effect occurred from 181 ± 25ms (mean ± sem) to 254 ± 22ms (mean ± sem). In the pre‐ and post-central gyri the multisensory integration occurred as a supra-additive non-linear neural response (AT > A+T) (Figure 3, for an exemplar LFP for AT multisensory integration). In the IFG the multisensory integration occurred as a sub-additive non-linear neural response (AT < A+T).

**Figure 3.**
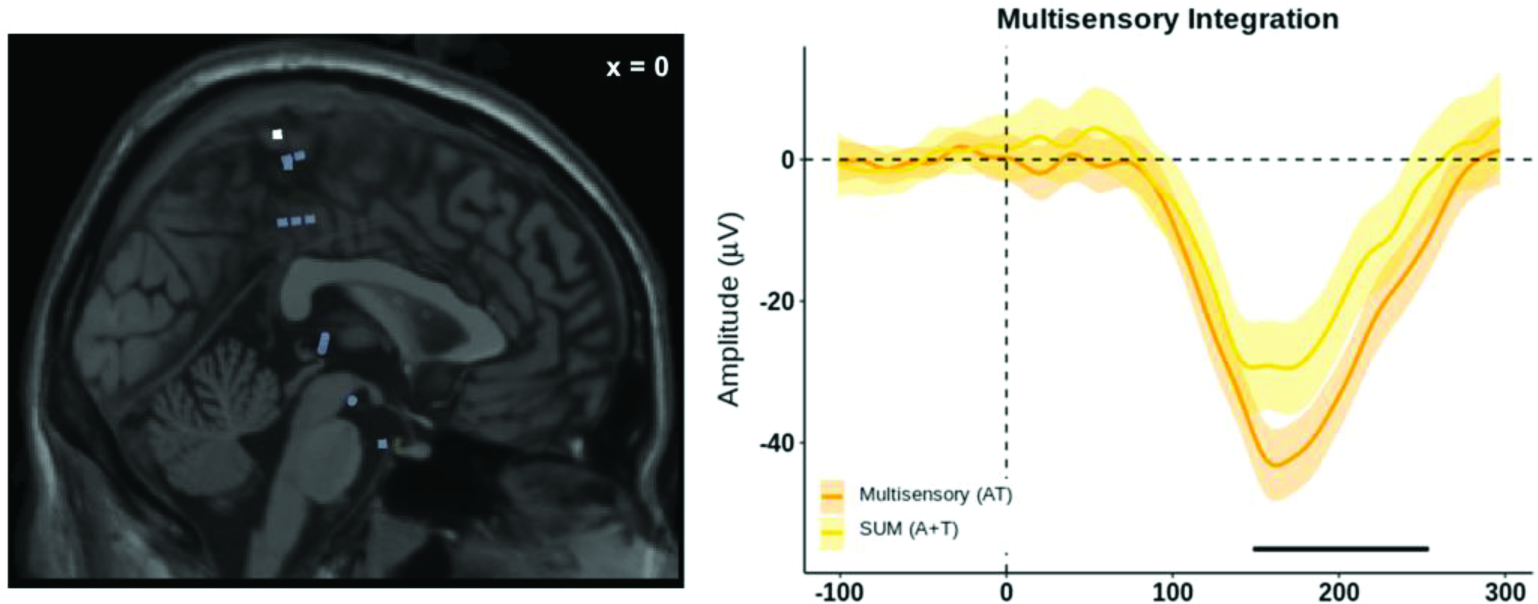
Exemplar LFP for AT multisensory integration. This electrode (highlighted in white in the left panel) showed a multisensory integration effect at 148-253ms after the stimulus onset and was located in the postcentral gyrus (PCG). The lines indicate (orange color for AT multisensory response & yellow color for the sum A + T) the average over trials; the shaded areas indicate the 95% C.I., and the black lines indicate the time period with a significant AT multisensory integration (p-value < 0.05, cluster-corrected).

### Location of the electrodes showing a PPS effect

Among the 20 electrodes showing AT multisensory integration, 6 electrodes (30% of AT multisensory electrodes, and 6% of active AT electrodes) showed a PPS effect. From the electrodes characterized as coding for PPS, 3 electrodes were located in the PHG (50% of the PPS electrodes), 2 electrodes were found in the PCG (34% of the PPS electrodes, and 1 electrodes in the IC (17% of the PPS electrodes; see Figure 2, Table 2 for details). Importantly, for all the locations where a PPS effect was observed, the response profile differed as a function of the distance from the trunk in such a way that PPS-dependent multisensory integration does not linearly decrease with distance, but is more similar to a step-function (see Figure 4, right panels).

**Table 2.** Number of significant electrodes showing AT multisensory integration and PPS effect, sorted by brain regions (p-value < 0.05, cluster-corrected).

| Region | Number of PPS/Multisensory Electrodes |
| --- | --- |
| Postcentral Gyrus | 2 / 7 |
| Superior Temporal Gyrus | 0 / 3 |
| Parahippocampal Gyrus | 3 / 3 |
| Inferior Frontal Gyrus | 0 / 1 |
| Precentral Gyrus | 0 / 1 |
| Insula | 1 / 1 |
| White matter | 0 / 4 |
| Total | 6 / 20 |

**Figure 4.**
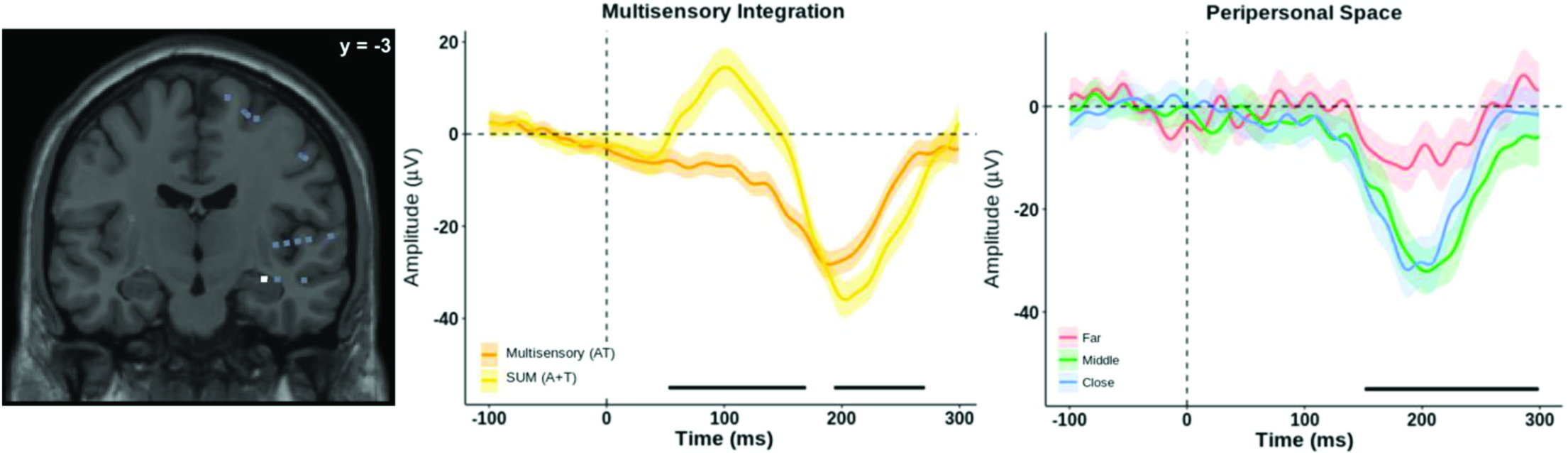
Exemplar LFP for AT multisensory integration & PPS effect. This electrode (highlighted in white in the left panel) showed a multisensory integration effect at 52-170ms and 193-270ms after stimulus onset and a PPS effect at 151-298ms after stimulus onset. The electrode was located in the parahippocampal gyrus. The lines indicate the average over trials; the shaded areas indicate the 95% C.I., and the black lines indicate the time period with a significant PPS effect (p-value < 0.05, cluster-corrected).

### Timing of the PPS effect

On average, across the 6 electrodes, the PPS effect occurred from 139ms ± 26ms (mean ± sem) to 226ms ± 31ms (mean ± sem) post-stimulus onset. Within the IC the effect occurred from 39ms to 129ms and from 141ms to 296ms. Within the PCG the PPS effect occurred from 44ms ± 3ms (mean ± sem) to 92ms ± 12ms (mean ± sem) and from 202ms ± 18ms (mean ± sem) to 265ms ± 7ms (mean ± sem). Within the PHG the PPS effect occurred from 193ms ± 23ms (mean ± sem) to 299ms ± 1ms (mean ± sem).

### Tactile habituation (control analysis)

To ascertain that the above-described PPS effect was not simply due to tactile habituation, we investigated if a ‘time-dependent’ effect (i.e., a significant difference on unisensory tactile responses as a function of the delay of tactile stimulation) was observed with the T stimulus alone. Among the 20 AT multisensory integration electrodes, 2 electrodes showed a possible anticipation/habituation effect in the tactile condition. These effects occurred within the PHG. Here, two distinct time periods showed a tactile habituation effect, on average the first time period occurred from 143ms ± 16ms (mean ± sem) to 213ms ± 35ms (mean ± sem) post-stimuli onset, and the second time period occurred from 188ms to 281 (only on one electrode – Figure S2). No other electrode showed a modulation in response to the T stimulus as a function of time.

These tactile habituation effects occurred during (at least partially) different time-points than the PPS effects, and the modulation was “linearly” dependent on the temporal order of the stimuli presentation. That is, the effect on the T condition showed a modulation of the LFPs for the 1^st^ vs. 2^nd^ administrated tactile stimulus (over both time significant time periods), and for the 2^nd^ vs. 3^rd^ (over the second significant time period) administrated tactile stimulus. This modulation pattern was different from what was observed for the PPS effect, where we see a difference, for instance, between the far and the middle/close distance (conceptually similar to a step-function, see Figure S2).

## Discussion

Intracranial EEG human recordings were performed in six patients suffering from pharmacoresistant epilepsy, who were presented with vibrotactile stimulation and concurrent sounds approaching their trunk in an effort to unravel the neurophysiological substrates of audio-tactile peripersonal space (PPS) surrounding the trunk. Crucially, and overlooked in most previous studies, PPS is defined as a multisensory spatial extent (e.g., Ferri et al., 2016; Graziano et al., 1997; Serino et al., 2015). Therefore, here we first identified brain responses that exhibited AT multisensory integration, indexed by non-linearity compared to the sum of the unisensory constituents of the multisensory stimuli (i.e., A+T vs. AT). Subsequently, within this subset of multisensory integration responses, we identified those that showed a modulation of the response as a function of distance from the body – that is, a PPS response. Broadly, results demonstrated that 104 (21%) of the 500 electrodes, implanted for clinical purposes, were responsive to the multisensory AT stimulation, with 104 electrodes (21%) and 78 (16%) being only responsive to T or the A unisensory stimulation, respectively. In addition, 19% (20 electrodes from the 104 electrodes showing an AT multisensory response) of these active electrodes specifically exhibited multisensory integration (defined as a non-linear summation of the response to AT stimuli, differently from the sum of A+T stimuli). These were located predominantly in the PCG and STG, but also in the IC and PHG. The AT multisensory response occurred, respectively, on average from 181ms, 100ms, 63ms, and 112ms. Among these 20 AT multisensory integration electrodes, 30% (6 electrodes) also showed a PPS effect – i.e., a non-linear modulation of the response to tactile stimuli as a function of the distance of the sounds from the body. Crucially, the spatial modulation of the responses did not linearly decrease with distance from the body (as it may be the case for responses related to tactile anticipation), but differentiated between the far vs. middle and/or near positions, suggestive of the presence of an electrophysiologically defined boundary between PPS and the far space (between 30cm and 100 cm from the body), in agreement with behavioral data in humans (Noel et al., 2015; Serino et al., 2015). The brain regions demonstrating a multisensory PPS effect were located most prominently in the PCG, but also within other cortical structures, namely the IC and the PHG (see Grivaz et al., 2017 for independent corroborative evidence). The PPS effect in those brain regions occurred, respectively, on average at 44ms, 39ms and 193ms. Hence, we present neurophysiological evidence, for the first time in humans, for the encoding of audio-tactile multisensory PPS in an extended cortical network. These human findings corroborate and extend those described in non-human primate studies, which demonstrated PPS processing mostly tested around the face and the hand in the parietal lobe (Duhamel et al., 1997, 1998; Graziano et al., 1997, 1999; Schlack et al., 2005) and fMRI studies in humans showing specific processing for stimuli presented close to the hand and face (parietal lobe, primary somatosensory cortex and insula; Grivaz et al., 2017). The present results describe neural mechanisms of the human *trunk* PPS that has been suggested to also be of particular importance for bodily self-consciousness (Noel et al., 2015; Serino et al., 2015; Blanke et al., 2015) and also reveal evidence for trunk PPS coding within the limbic system (i.e., PHG). In the following, we discuss our results with respect to multisensory integration, PPS and the conjunction between the two processes, in terms of brain location and timing of the effects.

### Location of audio-tactile multisensory integration

Our results corroborate and extend previous literature, by showing multisensory neural processing in response to audio-tactile multisensory pairings (vs. visuotactile in Avillac et al., 2007), to humans (vs. monkeys) and to LFP recordings (vs. single units). Research regarding multisensory integration - in particular, electrophysiological studies in non-primary sensory areas - have focused mostly on audio-visual and visuo-tactile integration. Much less is known about AT multisensory integration. Classically, multisensory integration has been considered to occur in higher-order temporal, parietal and occipital regions (Jones and Powell, 1970, more recently Quinn et al., 2014). However, this view has been challenged by studies, in both monkeys and humans alike, that provided evidence for early multisensory neural modulations (Lakatos et al., 2007; Schroeder and Foxe, 2002) occurring in regions traditionally considered purely unisensory cortices. Many of these modulatory effects in primary sensory areas have been demonstrated via somatosensory or visual effects in the primary auditory areas (Lakatos et al., 2007; Schroeder and Foxe, 2002). However, both fMRI and EEG experiments have highlighted the posterior superior temporal plane (Foxe et al., 2002; Murray et al., 2005), and not primary sensory areas, as brain regions implicated in veritable, overt or supra-threshold, audio-tactile multisensory integration. Other studies localized AT multisensory integration in the posterior parietal cortex, the somatosensory area SII and insula, rather than auditory association cortices (Gobbelé et al., 2003; Lütkenhöner et al., 2002; Renier et al., 2009). Thus, despite the fact that brain coverage in our study was limited by clinical purposes, the location of the electrodes showing stronger AT multisensory integration responses in our data corroborate and extend previous literature. That is, most of the electrodes demonstrating multisensory integration in our study were located in PCG, but also in the STG and IC. Despite, most of the multisensory electrodes being located in the PCG and thus anterior to the VIP region studied in monkeys (Avillac et al., 2007), this difference might be partially due to a different location of VIP in humans (more anterior and ventral) than in monkeys (Sereno and Huang, 2014).

### Timing of audio-tactile multisensory integration

A key advantage of recording intracranial LFPs as opposed to the BOLD response is that the former allows for indexing and characterizing time-resolved computations, in combination with high spatial resolution. In monkeys, AT multisensory integration has been reported to occur at early latencies (< 100ms) (e.g., Schroeder and Foxe, 2002; Schroeder et al., 2001, 2003). In humans, AT multisensory integration has been reported to occur both at early latencies (e.g., Murray et al., 2005) and at later latencies > 100ms (Lütkenhöner et al., 2002; Gobbelé et al., 2003). Late multisensory integration is supported by Quinn et al. (2014), who reported multisensory responses to visuo-tactile stimulation from iEEG recordings at latencies ranging from 145ms to 313ms poststimulus onset. Multisensory integration at later latency is also consistent with Valdés-Conroy et al., (2014), who performed an ERP study as a function of visual depth and reported a significant amplitude modulation in evoked responses within 150-200ms from stimulus onset. At first, our results show that the latency of the AT multisensory integration responses occurs on average between 151ms and 244ms, which is in agreement with later processes of multisensory integration. However, it is important to note that our results show three distinct temporal response patterns. That is, we also found an early AT multisensory integration occurring at ~60ms after the stimulus onset, within the IC and PHG, and a later effect occurring at ~100ms within the STG, followed by the even later PCG and IFG responses, suggesting that AT multisensory integration occurs over two (at least partially) distinct time periods, over more ventral (earlier effects) and more dorsal (later effect) regions (e.g., Reiner et al., 2009).

### Location of PPS effect

The present results extend the findings of a recent fMRI meta-analysis (Grivaz et al., 2017) study in humans, that aimed at identifying areas that consistently coded for PPS. This later study identified a portion of the PCG, including regions of area 1, 2 and 3b, as well as areas 5 and 40, as key PPS areas. Similarly, the present (and other) studies found PPS-like responses in the IC (Cappe et al., 2012; Schaefer et al., 2012) in humans. Thus, locating the bulk of multisensory PPS neural responses to the PCG (present study) is corroborated by recent functional neuroimaging literature on PPS processing.

We also found multisensory PPS responses in the IC and PHG. Regarding the IC, although it is a known area of multisensory convergence (e.g., Calvert et al., 2001; Renier et al., 2009; Rodgers et al., 2008), its direct electrophysiological implication in the multisensory mapping of PPS has not been previously established. However, the IC has been linked to changes in body ownership, self-identification and self-location after multisensory illusions, such as the rubber hand illusion (Blefari et al., 2017; Brozzoli et al., 2012; Grivaz et al., 2017; Tsakiris et al., 2007), the enfacement illusion (Apps et al., 2015), and the full body illusion (Ionta et al., 2014; Park et al., 2016, 2017). It is therefore considered a key area for the processing of multisensory cues underlying bodily self-consciousness (Blanke, 2012; Salomon et al., 2016; Seth, 2013). Notably, the illusions often used to study bodily self-consciousness rely on the manipulation of the spatiotemporal congruency of tactile cues on the body and visual cues from the external space, and have been shown to induce remapping of the PPS around the hand (Brozzoli et al., 2012), face (Maister et al., 2015) and trunk (Noel et al., 2015; Park et al., 2017). The present finding of PPS-related activity in the IC, hence, reinforces theoretical postulations (Blanke, 2012; Blanke et al., 2015; Serino et al., 2013) and psychophysical results (Noel et al., 2015; Salomon et al., 2017) highlighting the association between bodily self-consciousness and PPS representation. Lastly, the PHG has previously been categorized as a multisensory region (Tanabe et al., 2005), and a number of studies have suggested that the PHG is part of a network involved in processes relating to bodily self-consciousness (Tsakiris et al., 2007; Forget et al., 2015) as well as spatial navigation and self-location (Guterstam et al., 2015a, 2015b). The present data underline the PHGs involvement in multisensory PPS processing. This finding deserves further research concerning the potential role of this region as a hub between multisensory PPS processing, self-related processing, and its well-described role in memory and spatial navigation (Guterstam et al., 2015a, b). Interestingly, experimental alterations of bodily self-consciousness have been suggested to alter memory formation through activation of the hippocampal formation (Bergouignan et al., 2014). Thus, it may be proposed that the PHG serves as a gateway between the lower-level (multisensory) aspects of PPS and the implication of trunk-centered PPS in higher-order level of cognition such as egocentric processing (Canzoneri et al., 2016). A speculation that remains to be further tested (see Berthoz, 2000, for similar speculation).

### The timing of PPS effect

Evidence on the timing of AT PPS effect is currently lacking based both animal and humans studies. The only evidence concerning the timing of the PPS effect is provided by studies investigating visuo-tactile PPS. At first, our PPS results may appear somewhat late compared to previous electrophysiological findings of visuo-tactile PPS, as the present PPS responses occurred on average between 129ms and 226ms. For instance, evidence from single cell recording in monkeys (Avillac et al., 2007) show visuo-tactile, PPS-related responses occurring already at 68ms. In humans, Sambo and Forster, (2009) performed a visuo-tactile ERP study as a function of the spatial disparity of visuo-tactile stimuli in depth and observed a modulation in ERP amplitude at electrodes over the superior temporal lobe already at 100ms post-stimulus onset. Similarly, Cappe et al., (2012) showed an effect of distance for audio-visual stimuli starting at ~75ms. Although the average of our PPS effect occurred somewhat later compared to previous evidence, it is worth noting that our results show distinct response patterns. That is, the IC and PCG show a first response at ~40ms after stimulus onset, which is compatible with an early visuo-tactile PPS response. In addition, we also found a later response (~150ms after stimulus onset) in the IC, PCG, and PHG, in line with later responses observed in previous studies. These results suggest that the PPS responses (on the trunk) occur during at least two distinct time periods, and can occur simultaneously over different brain areas, largely overlapping with AT responses (see next section).

### Audio-tactile multisensory integration and PPS effect

Another finding worth highlighting regarding AT multisensory integration and the PPS effect is that both these processes appear to co-exist spatially and temporally (i.e., in the same electrodes and during similar time periods). AT multisensory integration is apparent on average from 151ms to 244ms post-stimuli onset, while PPS effect is discernable on average from 139ms to 226ms post-stimuli onset. If we look in greater detail into the different regions where both multisensory integration and PPS effect occurred, the timing of the two processes also overlapped. This observation hence provides – for the first time – evidence speaking in favor of a PPS representation, which is not yielded after a series of processes whereby multisensory integration occurs first and in different brain regions and then forwarded to different regions forming a PPS representation of the space surrounding a body. On the contrary, our results show for the first time that AT multisensory integration and PPS effect are concurrent during two time periods and across several brain regions. These electrophysiological findings suggest that PPS processing is based on a form of multisensory integration which, in addition, shows a clear spatial modulation of the response, in agreement with previous suggestions from neuropsychology (Farnè and Làdavas, 2000) or psychophysical (e.g., Noel et al., 2015; Serino et al., 2015) studies. Our data add critical insight by demonstrating that the representation of the space near (vs. far) from one’s body results from the processing of events/objects in the world involving the response of multisensory brain regions located in the PCG, but also in deeper and more medial areas (such as the IC and the PHG), likely harboring multisensory neurons with bodily-anchored and depth-restricted RFs (Fogassi et al., 1996; Graziano et al., 1997, Avillac et al., 2007; see Magosso, 2010 for a computational model of multisensory PPS representation).

### Conclusion

We describe electrophysiological responses linked to the multisensory integration of AT events and distinguished them from unimodal A or unimodal T responses as well as from simple AT summation responses. In addition, we showed that among these AT multisensory responses, LFPs from specific sites were modulated by the distance between the A and T component in a way that distinguishes near, peripersonal or bodily space (the trunk PPS) from spatial compartments far from the body. AT multisensory integration was first observed in the IC and during later phases in the STG, PCG, PHG, and IFG. A similar spatiotemporal response pattern was observed for the PPS effect but limited to IC, PHG, and PCG. Taken together, the present findings show – for the first time – that AT multisensory integration and PPS effect share common spatial and temporal processes, which go beyond previous single unit reports of multisensory integration in PPS neurons in area VIP (Avillac et al., 2007) to a number of other cortical areas, while also indexing multisensory PPS in humans, via audio-tactile, as opposed to visuotactile integration, and around the trunk as opposed to the hand or face.

## Acknowledgments

This work was supported by the Bertarelli Foundation and the Swiss National Science Foundation [grant number 320030_166643] to Olaf Blanke. Andrea Serino is supported by a Swiss National Foundation [grant PP00P3_163951 / 1].

---

**Figure S1.**
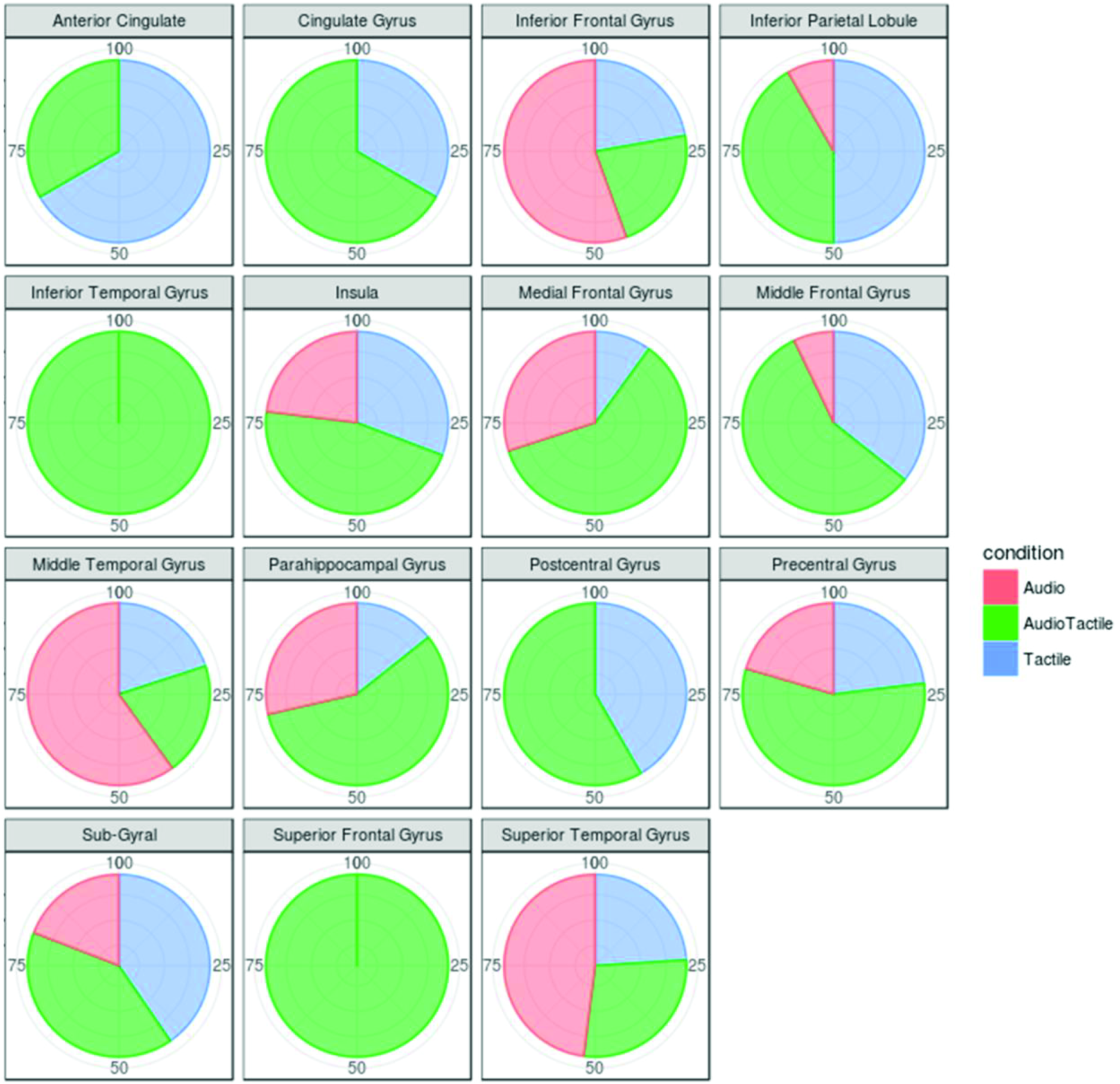
Electrode percentage per brain region. We show the percentage of response to the auditory, tactile and audio-tactile stimulation according to the brain regions.

**Figure S2.**
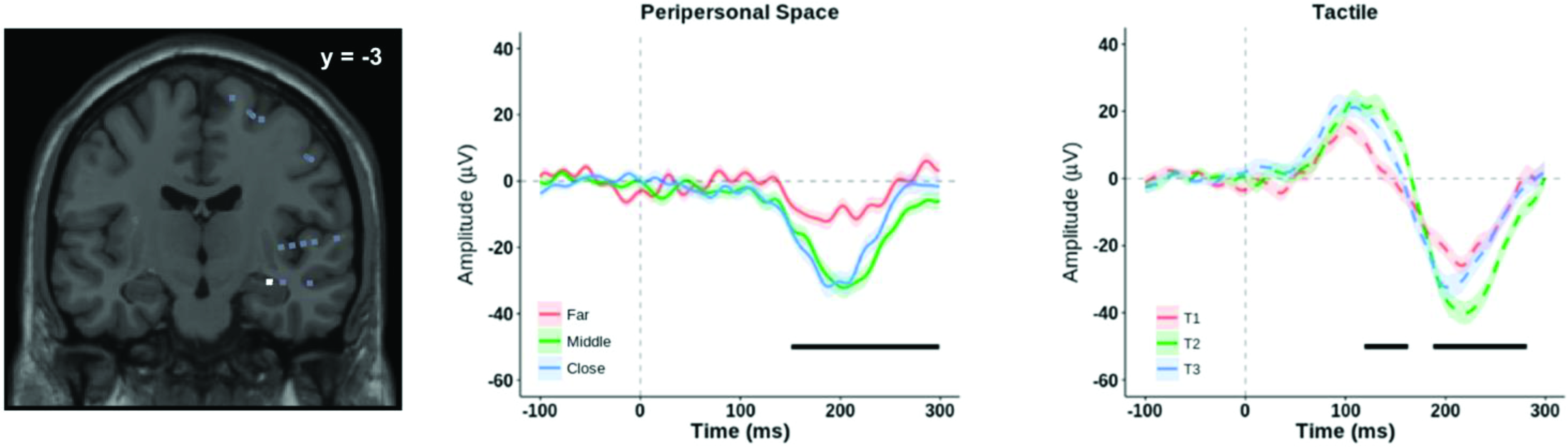
Exemplar LFP waveforms from one patient (control condition). This electrode (highlighted in white in the left panel) showed an audio-tactile multisensory integration (middle panel) effect at 52-170ms and 193-270ms after stimulus onset, a PPS effect (third panel from the left) at 151-298ms after stimulus onset and an effect was also observed for the unisensory tactile condition (fourth panel from the left) at 120-163ms and 188-281ms after the stimulus onset. The electrode was located in the parahippocampal gyrus. The lines indicate the average over trials; the shaded areas indicate the 95% C.I., and the black lines indicate the time period significant (p-value < 0.05, cluster-corrected).

**Supplementary Table S1.** Age, gender, handedness, epilepsy focus, heart rate, comorbidities, and medications (taken at the time of the recording) of the patients

| Patient | Age | Gender | Handedness | Epilepsy Focus | Medication |
| --- | --- | --- | --- | --- | --- |
| P-1 | 33 | Male | Left | Right post central epilepsy | No antiepileptic drugs |
| P-2 | 29 | Female | Left | Left parietal opercular epilepsy | Trileptal, Vimpat, Urbanyl |
| P-3 | 55 | Female | Right | Right temporal epilepsy | Trileptal |
| P-4 | 33 | Male | Right | Left temporal epilepsy | Carbamazepine |
| P-5 | 33 | Male | Right | Right post central epilepsy | Topiramate, Benzodiazepines |
| P-6 | 19 | Female | Right | Right temporal epilepsy | Valproate acid, Lamotrigine |

## References

Apps, M.A.J., Tajadura-Jiménez, A., Sereno, M., Blanke, O., and Tsakiris, M. (2015). Plasticity in unimodal and multimodal brain areas reflects multisensory changes in self-face identification. Cereb. Cortex N. Y. N 1991 25, 46‒55.

Avillac, M., Ben Hamed, S., and Duhamel, J.-R. (2007). Multisensory integration in the ventral intraparietal area of the macaque monkey. J. Neurosci. Off. J. Soc. Neurosci. 27, 1922‒1932.

Bergouignan, L., Nyberg, L., and Ehrsson, H.H. (2014). Out-of-body-induced hippocampal amnesia. Proc. Natl. Acad. Sci. U. S. A. 111, 4421‒4426.

Blanke, O. (2012). Multisensory brain mechanisms of bodily self-consciousness. Nat. Rev. Neurosci. 13, 556‒571.

Blanke, O., Slater, M., and Serino, A. (2015). Behavioral, Neural, and Computational Principles of Bodily Self-Consciousness. Neuron 88, 145‒166.

Blefari, M.L., Martuzzi, R., Salomon, R., Bello-Ruiz, J., Herbelin, B., Serino, A., and Blanke, O. (2017). Bilateral Rolandic operculum processing underlying heartbeat awareness reflects changes in bodily self-consciousness. Eur. J. Neurosci. 45, 1300‒1312.

Bremmer, F., Schlack, A., Shah, N.J., Zafiris, O., Kubischik, M., Hoffmann, K., Zilles, K., and Fink, G.R. (2001). Polymodal motion processing in posterior parietal and premotor cortex: a human fMRI study strongly implies equivalencies between humans and monkeys. Neuron 29, 287‒296.

Brozzoli, C., Gentile, G., Petkova, V.I., and Ehrsson, H.H. (2011). FMRI adaptation reveals a cortical mechanism for the coding of space near the hand. J. Neurosci. Off. J. Soc. Neurosci. 31, 9023‒9031.

Brozzoli, C., Gentile, G., and Ehrsson, H.H. (2012). That’s near my hand! Parietal and premotor coding of hand-centered space contributes to localization and self-attribution of the hand. J. Neurosci. Off. J. Soc. Neurosci. 32, 14573‒14582.

Brunet, D., Murray, M.M., and Michel, C.M. (2011). Spatiotemporal analysis of multichannel EEG: CARTOOL. Comput. Intell. Neurosci. 2011, 813‒870.

Calvert, G.A., Hansen, P.C., Iversen, S.D., and Brammer, M.J. (2001). Detection of audio-visual integration sites in humans by application of electrophysiological criteria to the BOLD effect. NeuroImage 14, 427‒438.

Canzoneri, E., Magosso, E., and Serino, A. (2012). Dynamic sounds capture the boundaries of peripersonal space representation in humans. PloS One 7, e44306.

Canzoneri, E., di Pellegrino, G., Herbelin, B., Blanke, O., and Serino, A. (2016). Conceptual processing is referenced to the experienced location of the self, not to the location of the physical body. Cognition 154, 182‒192.

Cappe, C., Thelen, A., Romei, V., Thut, G., and Murray, M.M. (2012). Looming signals reveal synergistic principles of multisensory integration. J. Neurosci. Off. J. Soc. Neurosci. 32, 1171‒1182.

Duhamel, J.R., Bremmer, F., Ben Hamed, S., and Graf, W. (1997). Spatial invariance of visual receptive fields in parietal cortex neurons. Nature 389, 845‒848.

Duhamel, J.R., Colby, C.L., and Goldberg, M.E. (1998). Ventral intraparietal area of the macaque: congruent visual and somatic response properties. J. Neurophysiol. 79, 126‒136.

Farnè, A., and Làdavas, E. (2000). Dynamic size-change of hand peripersonal space following tool use. Neuroreport 11, 1645‒1649.

Ferri, S., Pauwels, K., Rizzolatti, G., and Orban, G.A. (2016). Stereoscopically Observing Manipulative Actions. Cereb. Cortex N. Y. N 1991 26, 3591‒3610.

Fogassi, L., Gallese, V., Fadiga, L., Luppino, G., Matelli, M., and Rizzolatti, G. (1996). Coding of peripersonal space in inferior premotor cortex (area F4). J. Neurophysiol. 76, 141‒157.

Foxe, J.J., Wylie, G.R., Martinez, A., Schroeder, C.E., Javitt, D.C., Guilfoyle, D., Ritter, W., and Murray, M.M. (2002). Auditory-somatosensory multisensory processing in auditory association cortex: an fMRI study. J. Neurophysiol. 88, 540‒543.

Galli, G., Noel, J.P., Canzoneri, E., Blanke, O., and Serino, A. (2015). The wheelchair as a full-body tool extending the peripersonal space. Front. Psychol. 6, 639.

Gentile, G., Petkova, V.I., and Ehrsson, H.H. (2011). Integration of visual and tactile signals from the hand in the human brain: an FMRI study. J. Neurophysiol. 105, 910‒922.

Gentile, G., Guterstam, A., Brozzoli, C., and Ehrsson, H.H. (2013). Disintegration of multisensory signals from the real hand reduces default limb self-attribution: an fMRI study. J. Neurosci. Off. J. Soc. Neurosci. 33, 13350‒13366.

Gobbelé, R., Schürmann, M., Forss, N., Juottonen, K., Buchner, H., and Hari, R. (2003). Activation of the human posterior parietal and temporoparietal cortices during audiotactile interaction. NeuroImage 20, 503‒511.

Graziano, M.S.A., and Cooke, D.F. (2006). Parieto-frontal interactions, personal space, and defensive behavior. Neuropsychologia 44, 845‒859.

Graziano, M.S., Hu, X.T., and Gross, C.G. (1997). Visuospatial properties of ventral premotor cortex. J. Neurophysiol. 77, 2268‒2292.

Graziano, M.S., Reiss, L.A., and Gross, C.G. (1999). A neuronal representation of the location of nearby sounds. Nature 397, 428‒430.

Grivaz, P., Blanke, O., and Serino, A. (2017). Common and distinct brain regions processing multisensory bodily signals for peripersonal space and body ownership. NeuroImage 147, 602‒618.

Guterstam, A., Björnsdotter, M., Bergouignan, L., Gentile, G., Li, T.-Q., and Ehrsson, H.H. (2015a). Decoding illusory self-location from activity in the human hippocampus. Front. Hum. Neurosci. 9, 412.

Guterstam, A., Björnsdotter, M., Gentile, G., and Ehrsson, H.H. (2015b). Posterior cingulate cortex integrates the senses of self-location and body ownership. Curr. Biol. CB 25, 1416‒1425.

Holt, D.J., Cassidy, B.S., Yue, X., Rauch, S.L., Boeke, E.A., Nasr, S., Tootell, R.B.H., and Coombs, G. (2014). Neural correlates of personal space intrusion. J. Neurosci. Off. J. Soc. Neurosci. 34, 4123‒4134.

Huang, R.-S., Chen, C., Tran, A.T., Holstein, K.L., and Sereno, M.I. (2012). Mapping multisensory parietal face and body areas in humans. Proc. Natl. Acad. Sci. U. S. A. 109, 18114‒18119.

Ionta, S., Martuzzi, R., Salomon, R., and Blanke, O. (2014). The brain network reflecting bodily self-consciousness: a functional connectivity study. Soc. Cogn. Affect. Neurosci. 9, 1904‒1913.

Jones, E.G., and Powell, T.P. (1970). An anatomical study of converging sensory pathways within the cerebral cortex of the monkey. Brain J. Neurol. 93, 793‒820.

Làdavas, E., and Serino, A. (2008). Action-dependent plasticity in peripersonal space representations. Cogn. Neuropsychol. 25, 1099‒1113.

Lakatos, P., Chen, C.-M., O’Connell, M.N., Mills, A., and Schroeder, C.E. (2007). Neuronal oscillations and multisensory interaction in primary auditory cortex. Neuron 53, 279‒292.

Lancaster, J.L., Rainey, L.H., Summerlin, J.L., Freitas, C.S., Fox, P.T., Evans, A.C., Toga, A.W., and Mazziotta, J.C. (1997). Automated labeling of the human brain: a preliminary report on the development and evaluation of a forward-transform method. Hum. Brain Mapp. 5, 238‒242.

Lancaster, J.L., Woldorff, M.G., Parsons, L.M., Liotti, M., Freitas, C.S., Rainey, L., Kochunov, P.V., Nickerson, D., Mikiten, S.A., and Fox, P.T. (2000). Automated Talairach atlas labels for functional brain mapping. Hum. Brain Mapp. 10, 120‒131.

Leinonen, L., and Nyman, G. (1979a). II. Functional properties of cells in anterolateral part of area 7 associative face area of awake monkeys. Exp. Brain Res. 34, 321‒333.

Leinonen, L., and Nyman, G. (1979b). II. Functional properties of cells in anterolateral part of area 7 associative face area of awake monkeys. Exp. Brain Res. 34, 321‒333.

Lütkenhöner, B., Lammertmann, C., Simões, C., and Hari, R. (2002). Magnetoencephalographic correlates of audiotactile interaction. NeuroImage 15, 509‒522.

Magosso, E. (2010). Integrating information from vision and touch: a neural network modeling study. IEEE Trans. Inf. Technol. Biomed. Publ. IEEE Eng. Med. Biol. Soc. 14, 598‒612.

Maister, L., Cardini, F., Zamariola, G., Serino, A., and Tsakiris, M. (2015). Your place or mine: shared sensory experiences elicit a remapping of peripersonal space. Neuropsychologia 70, 455‒461.

Makin, T.R., Holmes, N.P., and Zohary, E. (2007). Is that near my hand? Multisensory representation of peripersonal space in human intraparietal sulcus. J. Neurosci. Off. J. Soc. Neurosci. 27, 731‒740.

Makin, T.R., Holmes, N.P., Brozzoli, C., Rossetti, Y., and Farnè, A. (2009). Coding of visual space during motor preparation: Approaching objects rapidly modulate corticospinal excitability in hand-centered coordinates. J. Neurosci. Off. J. Soc. Neurosci. 29, 11841‒11851.

Maravita, A., and Iriki, A. (2004). Tools for the body (schema). Trends Cogn. Sci. 8, 79‒86.

Maris, E., and Oostenveld, R. (2007). Nonparametric statistical testing of EEG‐ and MEG-data. J. Neurosci. Methods 164, 177‒190.

Murray, M.M., Molholm, S., Michel, C.M., Heslenfeld, D.J., Ritter, W., Javitt, D.C., Schroeder, C.E., and Foxe, J.J. (2005). Grabbing your ear: rapid auditory-somatosensory multisensory interactions in low-level sensory cortices are not constrained by stimulus alignment. Cereb. Cortex N. Y. N 1991 15, 963‒974.

Noel, J.-P., Grivaz, P., Marmaroli, P., Lissek, H., Blanke, O., and Serino, A. (2015). Full body action remapping of peripersonal space: the case of walking. Neuropsychologia 70, 375‒384.

Oostenveld, R., Fries, P., Maris, E., and Schoffelen, J.-M. (2011). FieldTrip: Open source software for advanced analysis of MEG, EEG, and invasive electrophysiological data. Comput. Intell. Neurosci. 2011, 156‒869.

Park, H.-D., Bernasconi, F., Bello-Ruiz, J., Pfeiffer, C., Salomon, R., and Blanke, O. (2016). Transient Modulations of Neural Responses to Heartbeats Covary with Bodily Self-Consciousness. J. Neurosci. Off. J. Soc. Neurosci. 36, 8453‒8460.

Park, H.-D., Bernasconi, F., Salomon, R., Tallon-Baudry, C., Spinelli, L., Seeck, M., Schaller, K., and Blanke, O. (2017). Neural Sources and Underlying Mechanisms of Neural Responses to Heartbeats, and their Role in Bodily Self-consciousness: An Intracranial EEG Study. Cereb. Cortex N. Y. N 1991 1‒14.

di Pellegrino, G., Làdavas, E., and Farné, A. (1997). Seeing where your hands are. Nature 388, 730.

Quinn, B.T., Carlson, C., Doyle, W., Cash, S.S., Devinsky, O., Spence, C., Halgren, E., and Thesen, T. (2014). Intracranial cortical responses during visual-tactile integration in humans. J. Neurosci. Off. J. Soc. Neurosci. 34, 171‒181.

Renier, L.A., Anurova, I., De Volder, A.G., Carlson, S., VanMeter, J., and Rauschecker, J.P. (2009). Multisensory integration of sounds and vibrotactile stimuli in processing streams for “what” and “where.” J. Neurosci. Off. J. Soc. Neurosci. 29, 10950‒10960.

Rizzolatti, G., Scandolara, C., Matelli, M., and Gentilucci, M. (1981). Afferent properties of periarcuate neurons in macaque monkeys. I. Somatosensory responses. Behav. Brain Res. 2, 125‒146.

Rizzolatti, G., Fadiga, L., Fogassi, L., and Gallese, V. (1997). The space around us. Science 277, 190‒191.

Rodgers, K.M., Benison, A.M., Klein, A., and Barth, D.S. (2008). Auditory, somatosensory, and multisensory insular cortex in the rat. Cereb. Cortex N. Y. N 1991 18, 2941‒2951.

Salomon, R., Ronchi, R., Dönz, J., Bello-Ruiz, J., Herbelin, B., Martet, R., Faivre, N., Schaller, K., and Blanke, O. (2016). The Insula Mediates Access to Awareness of Visual Stimuli Presented Synchronously to the Heartbeat. J. Neurosci. Off. J. Soc. Neurosci. 36, 5115‒5127.

Salomon, R., Noel, J.-P., Ltukowska, M., Faivre, N., Metzinger, T., Serino, A., and Blanke, O. (2017). Unconscious integration of multisensory bodily inputs in the peripersonal space shapes bodily self-consciousness. Cognition 166, 174‒183.

Sambo, C.F., and Forster, B. (2009). An ERP investigation on visuotactile interactions in peripersonal and extrapersonal space: evidence for the spatial rule. J. Cogn. Neurosci. 21, 1550‒1559.

Schaefer, M., Heinze, H.-J., and Rotte, M. (2012). Close to you: embodied simulation for peripersonal space in primary somatosensory cortex. PloS One 7, e42308.

Schlack, A., Sterbing-D’Angelo, S.J., Hartung, K., Hoffmann, K.-P., and Bremmer, F. (2005). Multisensory space representations in the macaque ventral intraparietal area. J. Neurosci. Off. J. Soc. Neurosci. 25, 4616‒4625.

Schroeder, C.E., and Foxe, J.J. (2002). The timing and laminar profile of converging inputs to multisensory areas of the macaque neocortex. Brain Res. Cogn. Brain Res. 14, 187‒198.

Schroeder, C.E., Lindsley, R.W., Specht, C., Marcovici, A., Smiley, J.F., and Javitt, D.C. (2001). Somatosensory input to auditory association cortex in the macaque monkey. J. Neurophysiol. 85, 1322‒1327.

Schroeder, C.E., Smiley, J., Fu, K.G., McGinnis, T., O’Connell, M.N., and Hackett, T.A. (2003). Anatomical mechanisms and functional implications of multisensory convergence in early cortical processing. Int. J. Psychophysiol. Off. J. Int. Organ. Psychophysiol. 50, 5‒17.

Sereno, M.I., and Huang, R.-S. (2006). A human parietal face area contains aligned head-centered visual and tactile maps. Nat. Neurosci. 9, 1337‒1343.

Sereno, M.I., and Huang, R.-S. (2014). Multisensory maps in parietal cortex. Curr. Opin. Neurobiol. 24, 39‒46.

Serino, A., Alsmith, A., Costantini, M., Mandrigin, A., Tajadura-Jimenez, A., and Lopez, C. (2013). Bodily ownership and self-location: components of bodily self-consciousness. Conscious. Cogn. 22, 1239‒1252.

Serino, A., Noel, J.-P., Galli, G., Canzoneri, E., Marmaroli, P., Lissek, H., and Blanke, O. (2015). Body part-centered and full body-centered peripersonal space representations. Sci. Rep. 5, 18603.

Seth, A.K. (2013). Interoceptive inference, emotion, and the embodied self. Trends Cogn. Sci. 17, 565‒573.

Spence, C., Pavani, F., Maravita, A., and Holmes, N. (2004). Multisensory contributions to the 3-D representation of visuotactile peripersonal space in humans: evidence from the crossmodal congruency task. J. Physiol. Paris 98, 171‒189.

Stein, B.E., and Stanford, T.R. (2008). Multisensory integration: current issues from the perspective of the single neuron. Nat. Rev. Neurosci. 9, 255‒266.

Stein, B.E., Meredith, M.A., and Wallace, M.T. (1993). The visually responsive neuron and beyond: multisensory integration in cat and monkey. Prog. Brain Res. 95, 79‒90.

Tadel, F., Baillet, S., Mosher, J.C., Pantazis, D., and Leahy, R.M. (2011). Brainstorm: a user-friendly application for MEG/EEG analysis. Comput. Intell. Neurosci. 2011, 879‒716.

Tanabe, H.C., Honda, M., and Sadato, N. (2005). Functionally segregated neural substrates for arbitrary audiovisual paired-association learning. J. Neurosci. Off. J. Soc. Neurosci. 25, 6409‒6418.

Teneggi, C., Canzoneri, E., di Pellegrino, G., and Serino, A. (2013). Social modulation of peripersonal space boundaries. Curr. Biol. CB 23, 406‒411.

Tsakiris, M., Hesse, M.D., Boy, C., Haggard, P., and Fink, G.R. (2007). Neural Signatures of Body Ownership: A Sensory Network for Bodily Self-Consciousness. Cereb. Cortex 17, 2235‒2244.

Tyll, S., Bonath, B., Schoenfeld, M.A., Heinze, H.-J., Ohl, F.W., and Noesselt, T. (2013). Neural basis of multisensory looming signals. NeuroImage 65, 13‒22.

Valdés-Conroy, B., Sebastián, M., Hinojosa, J.A., Román, F.J., and Santaniello, G. (2014). A close look into the near/far space division: a real-distance ERP study. Neuropsychologia 59, 27‒34.

